# A kernel-based method to calculate local field potentials from networks of spiking neurons

**DOI:** 10.1101/2020.03.29.014654

**Authors:** Bartosz Telenczuk, Maria Telenczuk, Alain Destexhe

## Abstract

**Background:** The local field potential (LFP) is usually calculated from current sources arising from transmembrane currents, in particular in asymmetric cellular morphologies such as pyramidal neurons.

**New method:** Here, we adopt a different point of view and relate the spiking of neurons to the LFP through efferent synaptic connections and provide a method to calculate LFPs.

**Results:** We show that the so-called unitary LFPs (uLFP) provide the key to such a calculation. We show experimental measurements and simulations of uLFPs in neocortex and hippocampus, for both excitatory and inhibitory neurons. We fit a “kernel” function to measurements of uLFPs, and we estimate its spatial and temporal spread by using simulations of morphologically detailed reconstructions of hippocampal pyramidal neurons. Assuming that LFPs are the sum of uLFPs generated by every neuron in the network, the LFP generated by excitatory and inhibitory neurons can be calculated by convolving the trains of action potentials with the kernels estimated from uLFPs. This provides a method to calculate the LFP from networks of spiking neurons, even for point neurons for which the LFP is not easily defined. We show examples of LFPs calculated from networks of point neurons and compare to the LFP calculated from synaptic currents.

**Conclusions:** The kernel-based method provides a practical way to calculate LFPs from networks of point neurons.

**Highlights:**

- We provide a method to estimate the LFP from spiking neurons
- This method is based on kernels, estimated from experimental data
- We show applications of this method to calculate the LFP from networks of spiking neurons
- We show that the kernel-based method is a low-pass filtered version of the LFP calculated from synaptic currents

## 1 Introduction

The local field potential (LFP) is the extracellular electric potential recorded using electrodes inserted in brain tissue. The LFP is thought to reflect mostly synaptic activity forming electric dipoles (Niedermeyer & Lopes da Silva, 1998; Nunez & Srinivasan, 2006), and can be well simulated using detailed morphologies (Bedard & Destexhe, 2012; Destexhe & Bedard, 2013; Lindén et al., 2014). However, it is not clear how to simulate LFPs from point neurons. In the present paper, we propose a method to calculate LFPs from point neurons, using experimentally-recorded LFP waveforms.

We focus on the unitary LFP (uLFP) which is the LFP generated by a single axon. The first investigations of uLFPs were done in hippocampal slices (Bazelot, Dinocourt, Cohen, & Miles, 2010; Glickfeld, Roberts, Somogyi, & Scanziani, 2009) and later in neocortex in vivo (B. Teleńczuk et al., 2017). In hippocampus, unitary LFPs were characterized in particular for inhibitory basket cells (Bazelot et al., 2010), which was convenient because the axon of a basket cell does not extend very far from the cell body (soma) and targets mostly the bodies and proximal dendrites of nearby pyramidal cells, and thus evokes postsynaptic currents clustered in space. In pyramidal neurons, however, efferent synapses target both basal and apical dendrites, and can extend far away from the cell, so in this case, the postsynaptic currents are rather scattered in space. This may be one of the reasons why the uLFP of pyramidal cells is much smaller in amplitude compared to that of inhibitory cells, as first suggested by Bazelot et al. (Bazelot et al., 2010).

This was taken one step further by Telenczuk et al. (B. Teleńczuk et al., 2017), who showed that uLFP can also be isolated from the neocortex *in vivo*, in human and monkey, where uLFPs could be extracted for both excitatory and inhibitory cells. Surprisingly, the two signals were of the same polarity despite being generated in principle by currents of opposite sign. Moreover, the excitatory uLFP was lagging behind the inhibitory uLFP. These properties let the authors to suggest that excitatory uLFPs may in fact be di-synaptic inhibitory uLFPs, which explains their polarity and timing relations. These properties were modeled by morphologically-detailed reconstructions of hippocampal neurons (M. Teleńczuk, Teleńczuk, & Destexhe, 2020), where it was shown that the weak uLFP of pyramidal cells is due to the scattering of their afferent synapses. Apical and basal excitatory synaptic currents produce LFPs of opposite sign, and there is therefore a significant amount of cancelling. However, inhibitory synapses in the somatic region always produce the same extracellular field, which explains why inhibitory uLFPs are of higher amplitude compared to excitatory uLFPs. Computational models fully support this explanation (M. Teleńczuk et al., 2020).

In the present paper, we take this another step further by showing that one can use the uLFPs as a powerful method to calculate LFPs. It was suggested previously (B. Teleńczuk et al., 2014, 2017; M. Teleńczuk et al., 2020) that one could use the measured uLFP as a basis to calculate LFP, but this was never attempted. We show here that this can be done by using templates from experimentally-recorded uLFPs, or from uLFPs calculated theoretically to estimate their spatial spread. The method consists of calculating the LFP of the network as a convolution of spiking activity with these uLFPs waveforms. The uLFPs thus constitute the “kernels” of such a convolution, hence the name “kernel-based method”. We illustrate this phenomenological method by calculating LFPs from networks of spiking neurons.

## 2 Materials and Methods

In the numerical test of the kernel-based method, we used network simulations of spiking neurons as described in previous papers (Brunel & Wang, 2003; Destexhe, 2009; Zerlaut, Chemla, Chavane, & Destexhe, 2018).

A first network (Brunel & Wang, 2003) consisted of 5,000 neurons, divided into 4,000 excitatory and 1,000 inhibitory cells, all described with the leaky integrate-and-fire model. The membrane time constant was of 20 ms and 10 ms for excitatory and inhibitory neurons, respectively, and the leak reversal potential was of -70 mV, the spike threshold was of -52 mV with a reset potential of -59 mV and an absolute refractory period of 2 ms for excitatory cells (1 ms for inhibitory cells). All cells were randomly connected with a connection probability *p* = 20%.

Synaptic currents were described as *I*_*syn*_(*t*) = *G*_*syn*_(*V* − *E*_*syn*_)*s*(*t*), where *G*_*syn*_ is the synaptic conductance, *E*_*syn*_ its reversal potential, and *s*(*t*) is a function describing the time course of synaptic currents and is described by the bi-exponential function

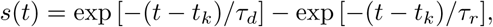

where *τ*_*r*_ is the rise time and *τ*_*d*_ the decay time of the postsynaptic conductance, and *t*_*k*_ is the time of the presynaptic spike. The reversal potential of excitatory (inhibitory) synaptic currents was 0 mV (−70 mV). The peak conductances were of 1 nS for excitatory synapses and 6 nS for inhibitory synapses. This network displays gamma-frequency (∼ 40 Hz) oscillations with sparse firing of all cell types (see (Brunel & Wang, 2003) for details).

In a second example, based on two previous papers (Destexhe, 2009; Zerlaut et al., 2018), we used networks of more complex integrate-and-fire models displaying spike-frequency adaptation, modeled by the Adaptive Exponential (AdEx) integrate-and-fire model (Brette & Gerstner, 2005). We considered a population of *N* = 10^4^ neurons randomly connected with a connection probability of *p* = 5%. We considered excitatory and inhibitory neurons, with 20% inhibitory neurons. The AdEx model permits to define two cell types, “regular-spiking” (RS) excitatory cells, displaying spike-frequency adaptation, and “fast spiking” (FS) inhibitory cells, with no adaptation. The dynamics of these neurons is given by the following equations:

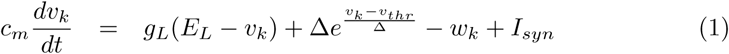

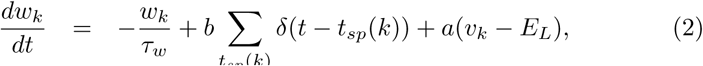

where *c*_*m*_ = 200 pF is the membrane capacitance, *v*_*k*_ is the voltage of neuron *k* and, whenever *v*_*k*_ *> v*_*thr*_ = −50 mV at time *t*_*sp*_(*k*), *v*_*k*_ is reset to the resting voltage *v*_*rest*_ = −65 mV and fixed to that value for a refractory time *T*_*refr*_ = 5 ms. The leak term *g*_*L*_ had a fixed conductance of *g*_*L*_ = 10 nS and the leakage reversal *E*_*L*_ was of -65 mV. The exponential term had a different strength for RS and FS cells, i.e. Δ = 2mV (Δ = 0.5mV) for excitatory (inhibitory) cells. Inhibitory neurons were modeled as fast spiking FS neurons with no adaptation (*a* = *b* = 0 for all inhibitory neurons) while excitatory regular spiking RS neurons had a lower level of excitability due to the presence of adaptation (while *b* varied in our simulations we fixed *a* = 4 nS and *τ*_*w*_ = 500 ms if not stated otherwise).

The synaptic current *I*_*syn*_ received by neuron *i* is the result of the spiking activity of all neurons *j* ∈ pre(*i*) pre-synaptic to neuron *i*. This current can be decomposed in the synaptic conductances evoked by excitatory E and inhibitory I pre-synaptic spikes

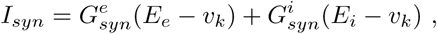

where *E*_*e*_ = 0mV (*E*_*i*_ =− 80mV) is the excitatory (inhibitory) reversal potential. Excitatory synaptic conductances were modeled by a decaying exponential function that sharply increases by a fixed amount *Q*_*E*_ at each pre-synaptic spike, i.e.:

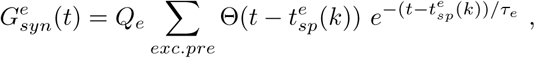

where Θ is the Heaviside function, *τ*_*e*_ = *τ*_*i*_ = 5ms is the characteristic decay time of excitatory and inhibitory synaptic conductances, and *Q*_*e*_ = 1 nS (*Q*_*i*_ = 5 nS) the excitatory (inhibitory) quantal conductance. Inhibitory synaptic conductances are modeled using the same equation with *e* → *i*. This network displays two different states according to the level of adaptation, *b* = 0.005 nA for asynchronous-irregular states, and *b* = 0.02 nA for Up/Down states (see (Zerlaut et al., 2018) for details).

In some simulations, we compared the kernel method to a classic method to compute local field potentials from the synaptic currents. In this case, the extracellular potential *V*_*e*_ at a position 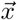 was computed as:

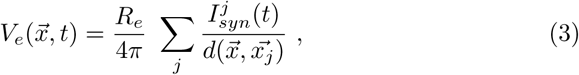

where *R*_*e*_ = 230*cm* is the extracellular resistivity, 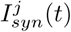 (*t*) is the synaptic current of neuron *j* as defined above, and 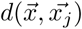 is the distance between 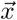 and the position of neuron *j*, 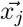.

All simulations were done using the NEURON Simulation environment (Hines & Carnevale, 1997) or the BRIAN simulator (Goodman & Brette, 2009). Program codes for the network models are available from the original papers (Brunel & Wang, 2003; Destexhe, 2009; Zerlaut et al., 2018). The program code of the kernel method is available open-access (B. Teleńczuk, Teleńczuk, & Destexhe, 2020) (using the hoc language of NEURON, as well as in python 3).

## 3 Results

We start by showing the essential properties of uLFPs as recorded experimentally, and then consider a method to generate LFPs based on those measurements. We also show the results from a detailed biophysical model of uLFPs, which we use to infer the depth-dependence of the model of uLFPs. Finally, we illustrate the method by calculating LFPs from networks of spiking point neurons.

### 3.1 Unitary local field potentials

Figure 1 illustrates the properties of uLFPs. The uLFP is generated by a single axon, where all efferent synapses of the axon collateral (schematized in Fig. 1A) will generate a small field due to the postsynaptic current, and the ensemble of these small fields constitutes the uLFP. One property of the uLFP is that recordings made at different distances from the soma will peak at different times, because of the speed of action potential propagation along the axon (Fig. 1B). Thus, the peak time of the uLFP as a function of distance is expected to show a linear increase, as schematized in Fig. 1C, where the slope is the axon propagation speed. In addition, the uLFP should also display a peak decreasing with distance, as expected for electrodes located at increasing distances from the soma (Fig. 1D).

**Figure 1:**
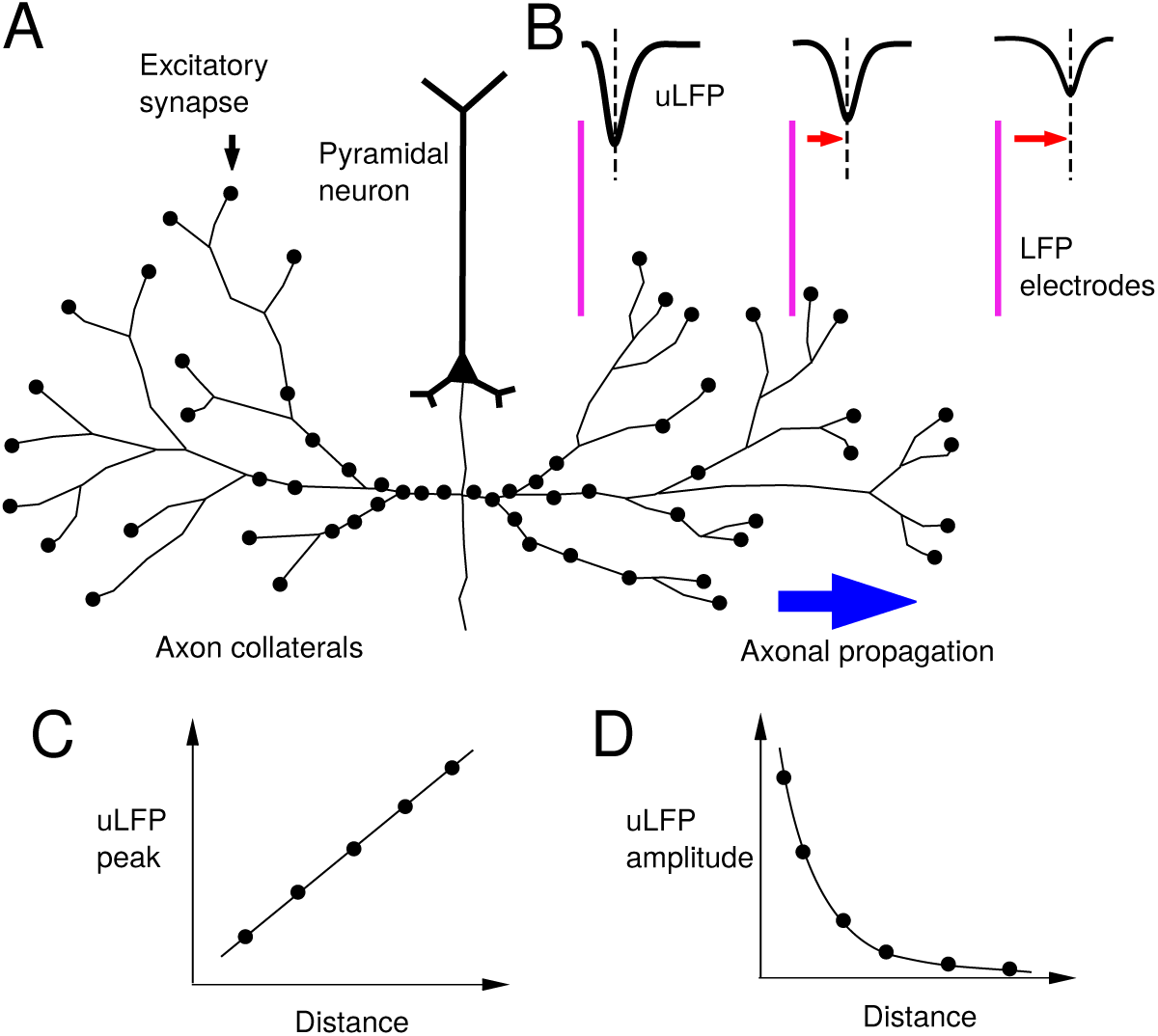
Scheme of the genesis of unitary LFPs. A. Scheme of the axonal arborization of a pyramidal cell, where the axon collaterals extend laterally and contact other neurons in the vicinity of the cell. Black dots indicate excitatory synapses made my the axon on different neurons. B. Scheme of 3 LFP electrodes (violet) located at different distances from the soma. The uLFP recorded by each electrode is progressively delayed (red arrows), due to the limited speed propagation along the axon (blue arrow). The amplitude is also progressively lower due to increasing distances from the sources. C. Scheme of the peak time of the uLFP as a function of distance, which reflects axonal propagation. D. Scheme of the decrease of uLFP amplitude as a function of distance.

These properties were found in human recordings by a previous study (B. Teleńcz et al., 2017), as summarized in Fig. 2. Recordings were made using Utah arrays inserted in temporal cortex, leading to LFP and unit recordings (Fig. 1A). Telenczuk et al. (B. Teleńczuk et al., 2017) used a whitening method to extract the relation between unit spikes and the LFP. The properties of this relation reminds those of the uLFP. First, the presumed uLFP peak amplitude decreases with distance, with an exponentially decaying function with a space constant around 200 *µ*m, consistent with other estimates (Katzner et al., 2009). Second, the uLFP peak scaled linearly with distance, with an estimated speed of 200 mm/sec (Fig. 2C), which is consistent with the action potential speed along axon unmyelinated fibers. These properties were also found for a second human subject recorded similarly (Fig. 2D). Very similar uLFP waveforms were obtained from monkey motor cortex, which was recorded with similar Utah arrays. The details of this analysis can be found in ref. (B. Teleńczuk et al., 2017).

**Figure 2:**
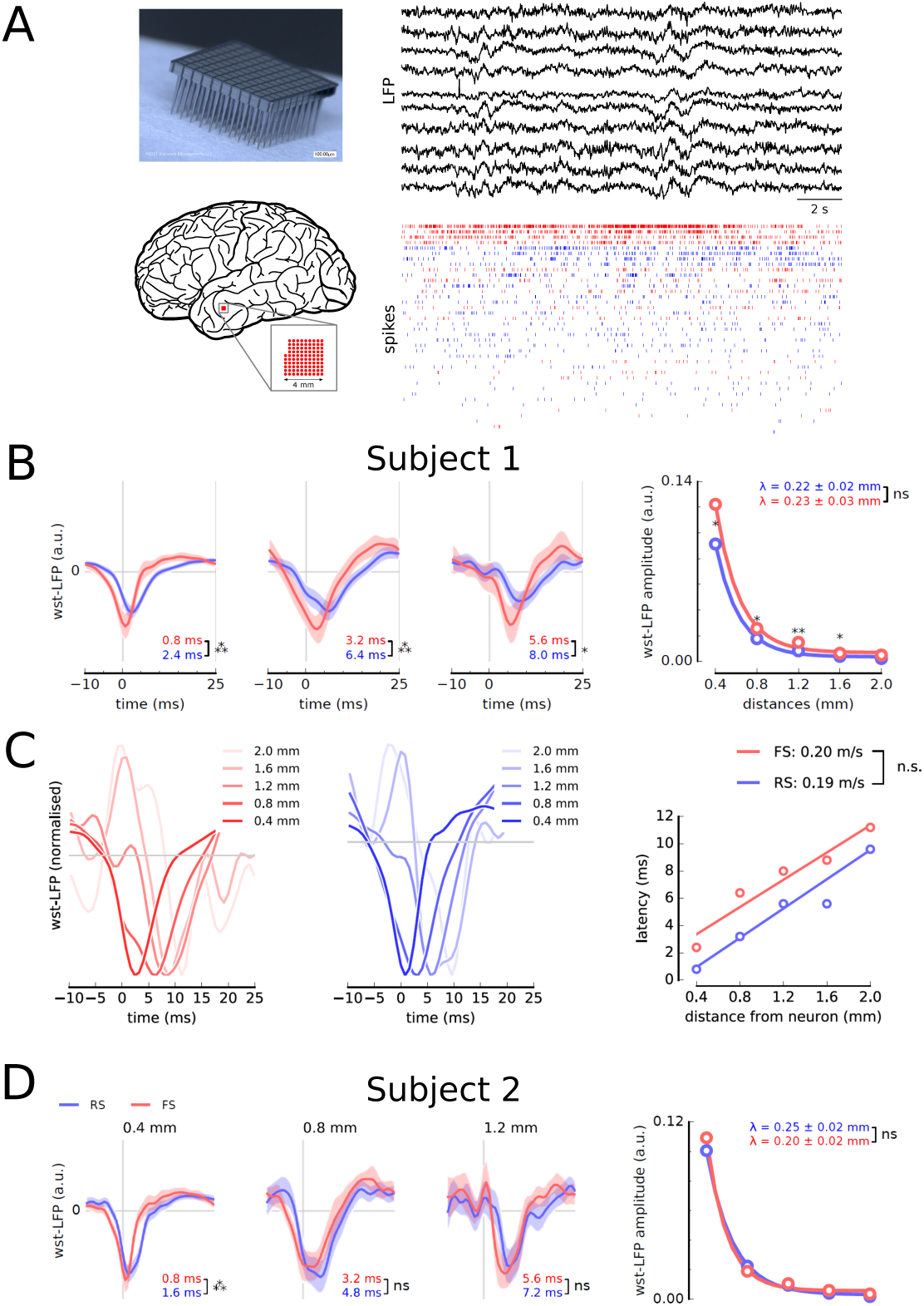
Experimental measurements of unitary LFPs in human. A. Utah-array (top left) recording in human temporal cortex (bottom left), of LFPs and units (right traces). 10 example LFP traces are shown, along with spike-sorted units, which are represented from top to bottom as a decreasing function of their mean firing rate. Presumed excitatory (RS, blue) and inhibitory (FS, red) cells are indicated. B. Unitary LFPs for RS (blue) and FS (red) neurons at different electrode distances. The rightmost graph shows the uLFP amplitude as a function of distance. C. Traveling of uLFPs (left graphs). The uLFP peak travels at a speed close to 200 mm/sec (right), consistent with axonal propagation. D. Results obtained from a second subject, in agreement with B. Modified from (B. Teleńczuk et al., 2017).

Another important information is the respective excitatory and inhibitory contributions to LFPs. It can be seen from Fig. 2 that the uLFP from excitatory or inhibitory cells are of the same polarity (negative in this case). However, the synaptic currents generating these uLFPs are of opposite sign, so they should lead to opposite polarities. It was proposed (B. Teleńczuk et al., 2017) that this is due to the fact that excitatory uLFPs are of low amplitude compared to inhibitory uLFPs, and the field evoked by excitatory cells is actually dominated by inhibitory currents, occurring di-synaptically through the recruitment of interneurons. This explanation was consistent with hippocampal recordings, where the excitatory uLFPs were of very small amplitude and blocked by GABA_*A*_ antagonists (Bazelot et al., 2010). This issue was tested in a recent biophysical model (M. Teleńczuk et al., 2020), which reconstructed uLFPs in the hippocampus for excitatory and inhibitory synapses. By using 1000 morphologically-reconstructed hippocampal pyramidal neurons (Fig. 3A), and locating synapses in different regions of the cells (Fig. 3B-C), the model generated uLFPs that were recorded at different positions around the cell (Fig. 3D-E). As suggested before, the model confirmed that inhibitory uLFPs (Fig. 3D) were of larger amplitude compared to excitatory uLFPs (Fig. 3E; see Overlay).

**Figure 3:**
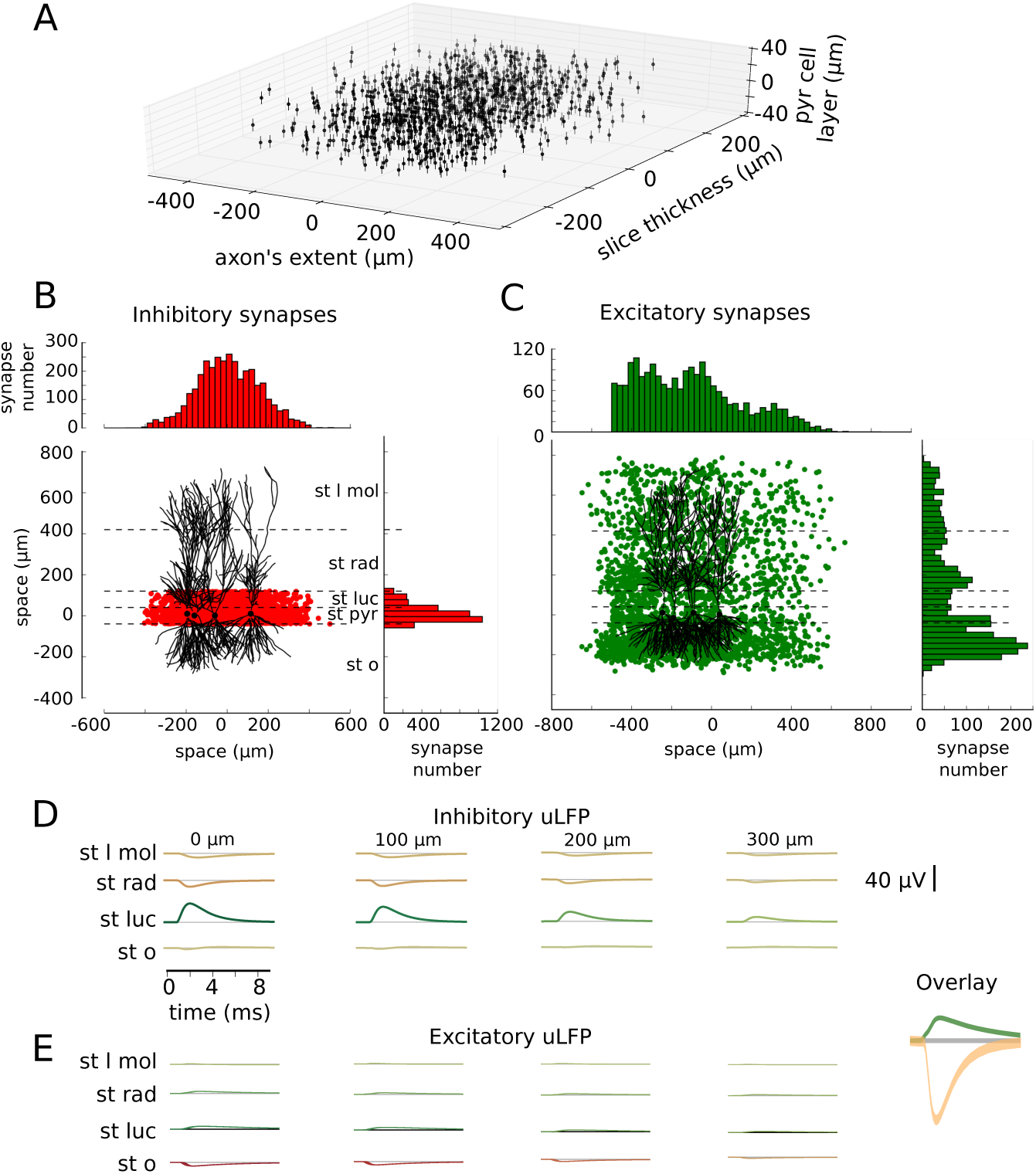
Detailed biophysical model of uLFPs in the hippocampus. A. Relative position of 1000 morphologically-reconstructed hippocampal CA3 pyramidal cells. B. Distribution of inhibitory synapses from basket cells, mainly targeting the somatic region of pyramidal cells. C. Distribution of excitatory synapses, mainly targeting apical and basal dendrites. D. Simulated uLFP from inhibitory neurons at different distances from the cell (resp. 0, 100, 200, 300 *µ*m, from left to right). E. Simulated uLFPs from excitatory neurons. There was a lot of cancelling for excitatory uLFPs, resulting in lower uLFP amplitudes compared to inhibitory uLFPs (Overlay, uLFPs indicated for stratum radiatum; 10x amplitude magnification). Modified from (M. Teleńczuk et al., 2020).

For this reason, in the following, we will fit kernel templates only to inhibitory uLFPs measured experimentally, and use the model to estimate kernels for excitatory uLFPs. Note that the procedure and examples shown here concern cerebral cortex, where the uLFP kernels were measured from experimental data (B. Teleńczuk et al., 2017), but the exact same approach can be followed for any brain structure where the uLFP kernels are measured experimentally.

### 3.2 Fitting kernels to inhibitory uLFPs

In this section, we fit a template kernel function to inhibitory uLFPs extracted from experimental data in cerebral cortex. To do this, we note that the uLFPs always have an approximately symmetric shape with similar rise and decay phases, which we can fit by the following Gaussian kernel at position *x* and time *t*:

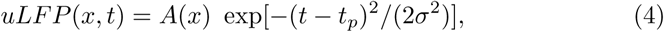

where *A* is an amplitude constant (which can be negative), *σ* is the standard deviation in time, and *t*_*p*_ is the peak time of the uLFP. The latter is given by

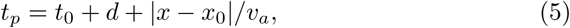

where *t*_0_ is the time of the spike of the cell, |*x* − *x*_0_| is the distance between cell and electrode, *d* is a constant delay, and *v*_*a*_ is the axonal speed. We use the value of *v*_*a*_ = 200 mm/s, estimated from human uLFP recordings (B. Teleńczuk et al., 2017).

To model the observed near-exponential amplitude decay with distance (Fig. 2B,D, rightmost graphs), the following expression can be used for *A*(*x*):

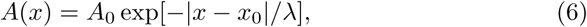

where *A*_0_ is the maximal amplitude, and |*x* − *x*_0_| is the distance between the electrode (*x*) and the position of the cell (*x*_0_), and *λ* is the space constant of the decay. From human uLFP data, *λ* was consistently found around 200-250 *µ*m (B. Teleńczuk et al., 2017).

The template function given by Eq. 4 can be fit simultaneously to sets of recorded LFPs, such as that of Fig. 2B. Figure 4 shows the result of such a fitting, for inhibitory uLFPs. The Gaussian kernel function with a negative amplitude, and constant standard deviation, could simultaneously fit all measured uLFPs (Fig. 3A). Note that we did not attempt to capture the slow positive component which is present in some of the uLFPs. Better fits can be obtained by letting the amplitudes and standard deviation as free parameters (Fig. 3B), but this type of parameterization is unconstrained, and will not be used in the following.

**Figure 4:**
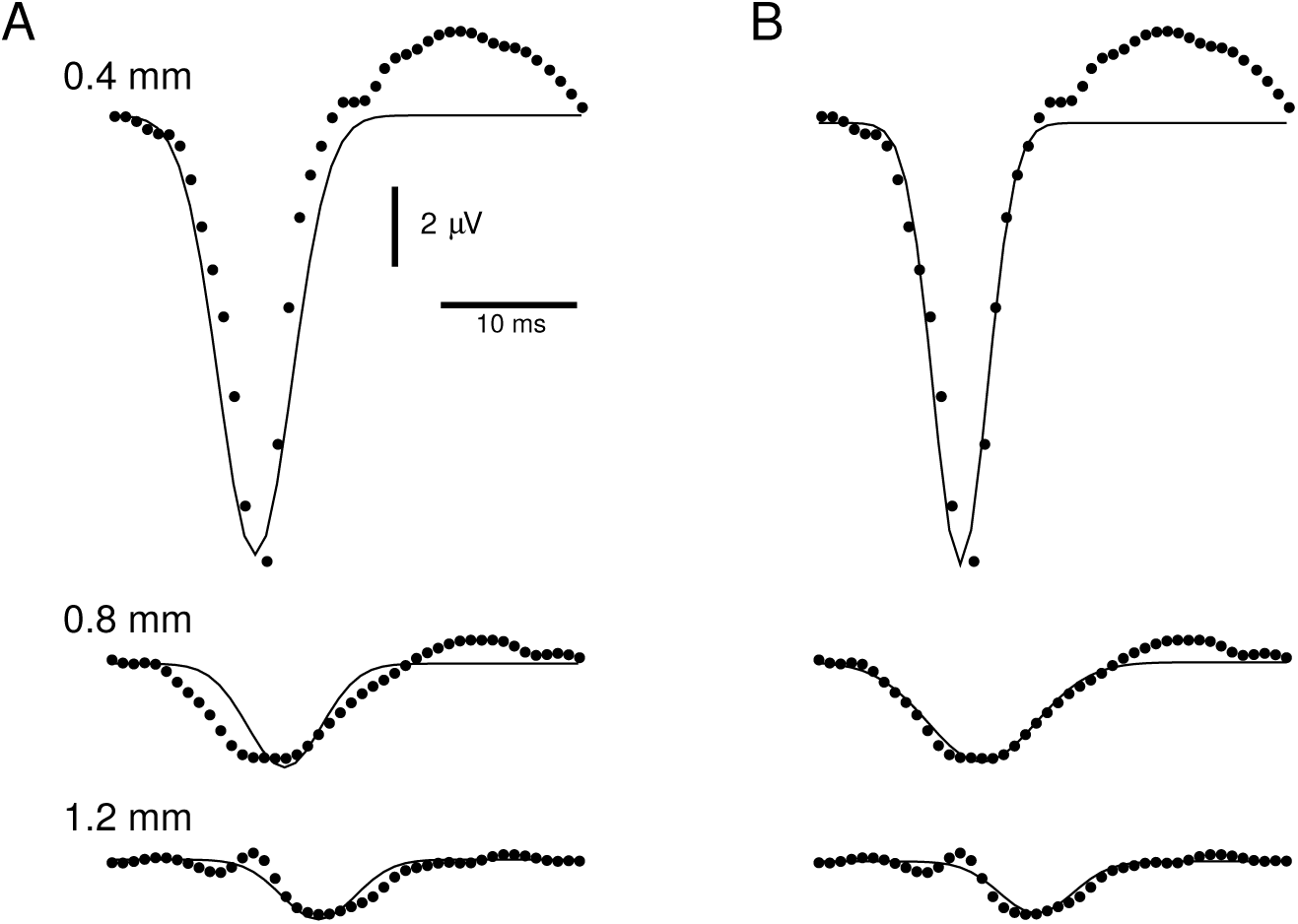
Fitting of Gaussian kernels to inhibitory uLFPs. A. Simultaneous fit of the same Gaussian template to inhibitory uLFPs measured experimentally (dots) at three different distances *x*. The template had constant standard deviation *σ*, and the amplitude was given by an exponentially-decaying function of distance (continuous curves; parameters: *v*_*a*_=166 mm/s, *d*=10.4 ms, *A*_0_=-3.4 *µ*V, *λ*= 0.34 mm, *σ*=2.1 ms). B. Similar fit using Gaussian templates with unconstrained parameters (amplitudes of -11, -2.5 and -1.4 *µ*V, and standard deviations of 1.99, 3.95 and 2.7 ms, respectively from top to bottom).

### 3.3 Calculating excitatory uLFPs

The fitting of the kernels provided in the previous section is enough to calculate the uLFP contribution of a given inhibitory cell at any point in space and time in the vicinity of the cell. However, as mentioned above, it is difficult to directly observe the uLFP of excitatory cells because of its low amplitude (Bazelot et al., 2010). In this section, we provide an estimate of the excitatory uLFP, based on numerical simulations. We use a biophysical model proposed previously (M. Teleńczuk et al., 2020), summarized in Fig. 3. This biophysical model confirmed that the simulated excitatory uLFP is indeed smaller compared to the inhibitory uLFP (compare Fig. 3D-E), although the number of synapses involved in calculating excitatory uLFPs was much larger compared to inhibitory synapses. The low amplitude of excitatory uLFP resulted from a partial cancellation of apical and basal synaptic currents, which produce dipoles of opposite sign (M. Teleńczuk et al., 2020). In the Overlay of Fig. 3 (from *stratum radiatum*), it can be seen that the excitatory uLFP is not only of smaller amplitude, but also generally has slower kinetics, presumably because of the distal dendritic contributions and associated dendritic filtering (Pettersen & Einevoll, 2008). Thus, for the kernel, we assumed a slower decay time for excitatory uLFP, which we estimated as about 1.5 times the decay of inhibitory uLFPs. More precise measurements, when available, should be used to adjust this number.

To estimate the relative amplitudes of excitatory and inhibitory uLFPs, we use the depth profile of uLFP, as shown in Fig. 5. It is apparent that the major contribution of inhibitory uLFPs will be around the soma (*stratum pyramidale*), with the two main poles reversing around 200 *µ*m depth (*stratum radiatum*), reversing again around 600 *µ*m. Excitatory uLFPs are also maximal around the soma, reverse around -100 *µ*m, but stay of low amplitude all through the layers. To simplify, we have reported the absolute and relative uLFP amplitudes at different depth in Table 1.

**Table 1:**
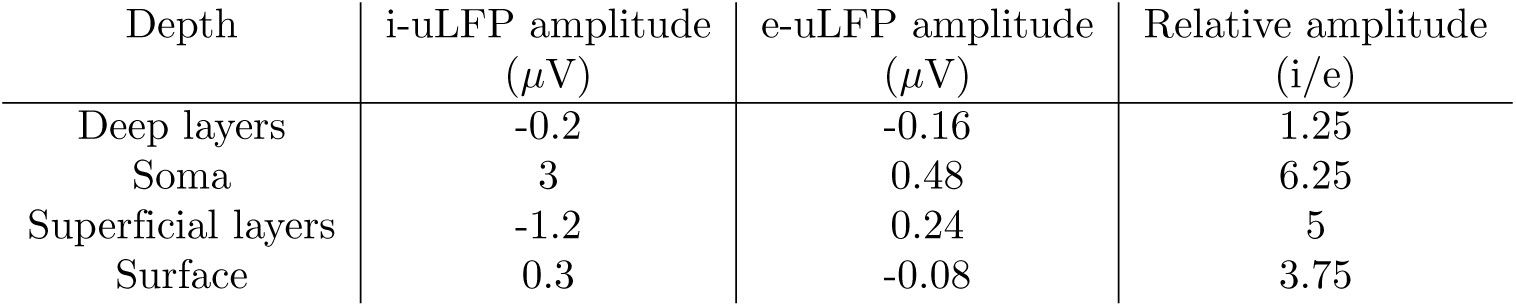
Absolute and relative model uLFP amplitudes at different depth in hippocampus. The uLFP amplitudes are indicated for a position near the soma in X,Y, and for different depths in Z direction. The different depths indicated correspond to -400 *µ*m (Deep layers), 0 (Soma), 400 *µ*m (Superficial layers) and 800 *µ*m (Surface).

**Figure 5:**
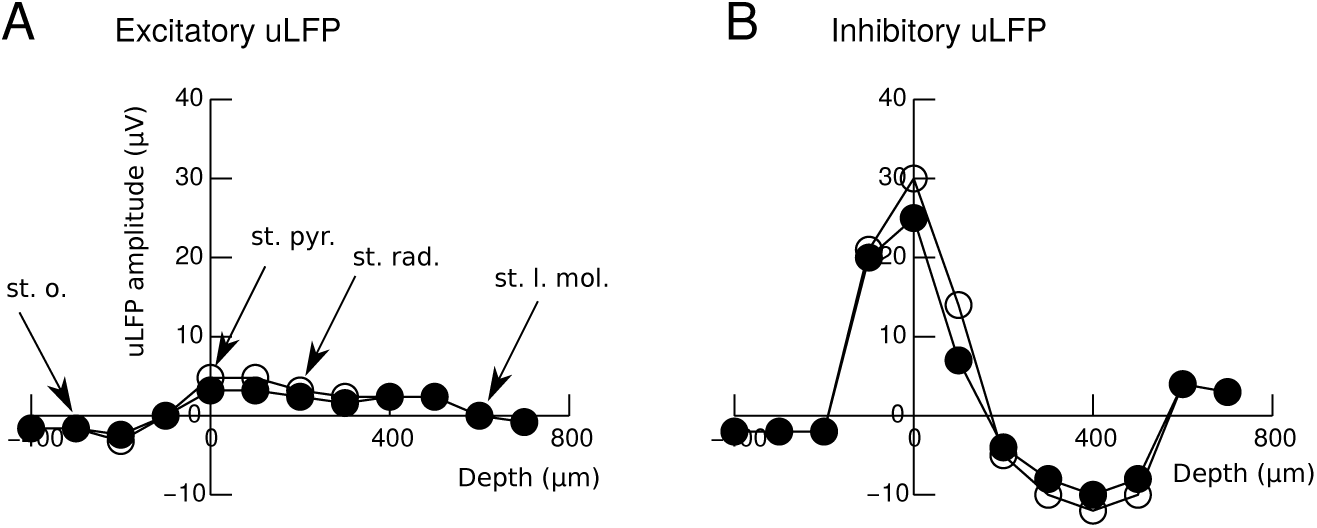
Depth profile of model uLFP peak amplitude in hippocampus. A. Peak uLFP amplitude as a function of depth (with zero in stratum pyramidale, as in Fig. 3B). B. Depth profile of peak amplitudes for inhibitory uLFPs. Open and filled circles indicate the position of 0 and 200 *µ*m (same scale in Fig. 3B).

With respect to the fitting of inhibitory uLFPs in the previous section, we obtained an amplitude of about -3.4 *µ*V and a width of 2.1 ms (Fig. 4A). Given that the corresponding recordings (B. Teleńczuk et al., 2017) were obtained in superficial layers, we can assume that it corresponds to superficial layers in Table 1. Accordingly, we estimate that excitatory uLFPs would have an amplitude of about 0.7 *µ*V and a width of 3.15 ms. We consider a practical application of these kernel templates to calculate LFPs in the next section.

### 3.4 Examples of LFPs calculated from network simulations

To calculate LFPs from network simulations, we will convolve the spikes of the network with the uLFP kernels, according to the formula:

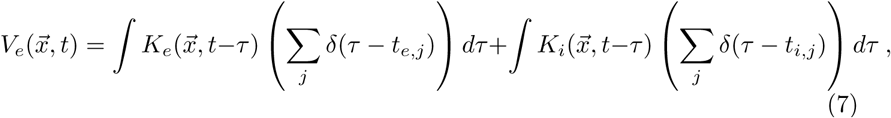

where 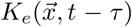 and 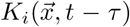 are the excitatory and inhibitory uLFP kernels derived above, respectively, while {*t*_*e,j*_} and {*t*_*i,j*_}are the spike times of excitatory and inhibitory neurons. This can also be expressed as a direct sum of the kernels:

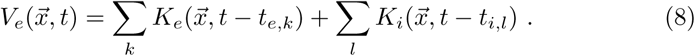

For convenience, we will use the LFP kernels estimated from human recordings using the Gaussian template (Eq. 4), as in Fig. 4.

Figure 6 shows an example of LFP generated using this kernel method applied to a network of spiking point neurons. The network was taken from a previous study, modeling gamma oscillations in networks of excitatory and inhibitory integrate-and-fire neurons (Brunel & Wang, 2003). As seen from the raters of spiking activity (Fig. 6A, top), the network displayed mostly irregular behavior, but at closer scrutiny (Fig. 6B, top), signs of loosely synchronized oscillatory activity can be seen. When calculating the LFP from this network (Fig. 6, bottom traces) clearly reveals the gamma oscillation in the LFP. The LFP was calculated using the templates estimated above, and using amplitudes as in Table 1 to simulate the LFP in different layers. One can see that the LFP is largest at the level of the soma, while surface and deep layers display lower amplitudes. The gamma oscillation also reverses in polarity above and below the soma layer. Figure 7 shows the LFP calculated at the level of the soma, but at different lateral distances from the center of the network plane. Note that these different traces are not scaled versions of the same trace, because each neuron contributes individually according to its distance to the electrode (see Eqs. 4-6). This shows that the kernel method reproduces the typical attenuation with distance as expected.

**Figure 6:**
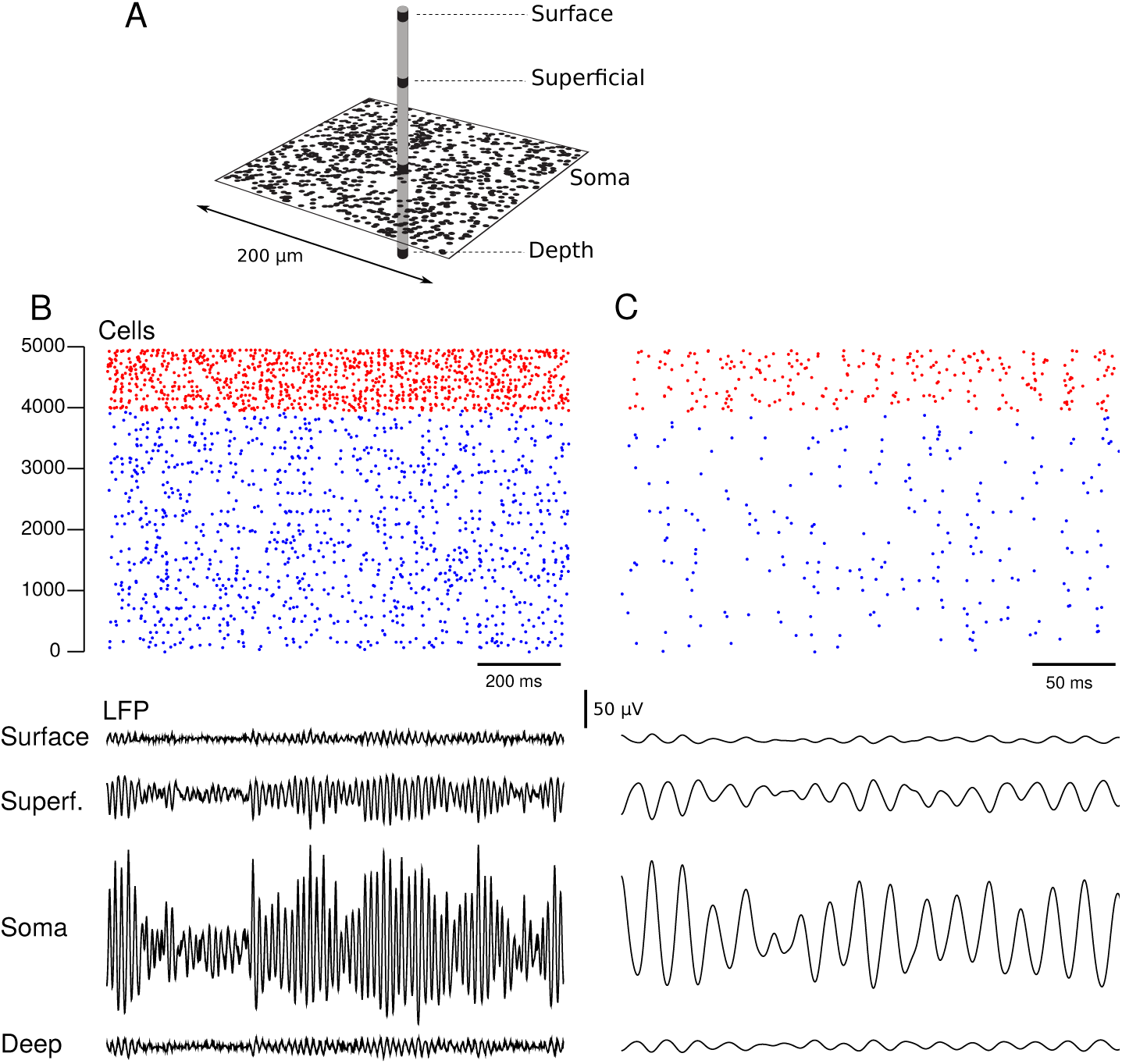
Example of LFP calculated from networks of point neurons exhibiting gamma oscillations. A. Scheme of the placement of cells and electrodes. Neurons were distributed randomly in a plane of 200 *µ*m size, and the electrodes were placed perpendicular to the plane, as indicated. B. Simulations of gamma oscillations in randomly-connected networks of excitatory and inhibitory neurons. The top graphs display the raster of spiking activity in the network. The network had 5,000 neurons, 4,000 excitatory (blue) and 1,000 inhibitory (red). The network models the genesis of gamma oscillations by recurrent excitatory and inhibitory interactions among integrate-and-fire neurons (Brunel & Wang, 2003). The bottom curves show the LFP calculated using the kernel method. From top to bottom: surface LFP, LFP from superficial layers, LFP at the level of the soma, and LFP in deep layers as schematized in A. The corresponding uLFP amplitudes were taken from Table 1, and the kernel were estimated as in Fig. 4. C. Same simulation at higher temporal resolution.

**Figure 7:**
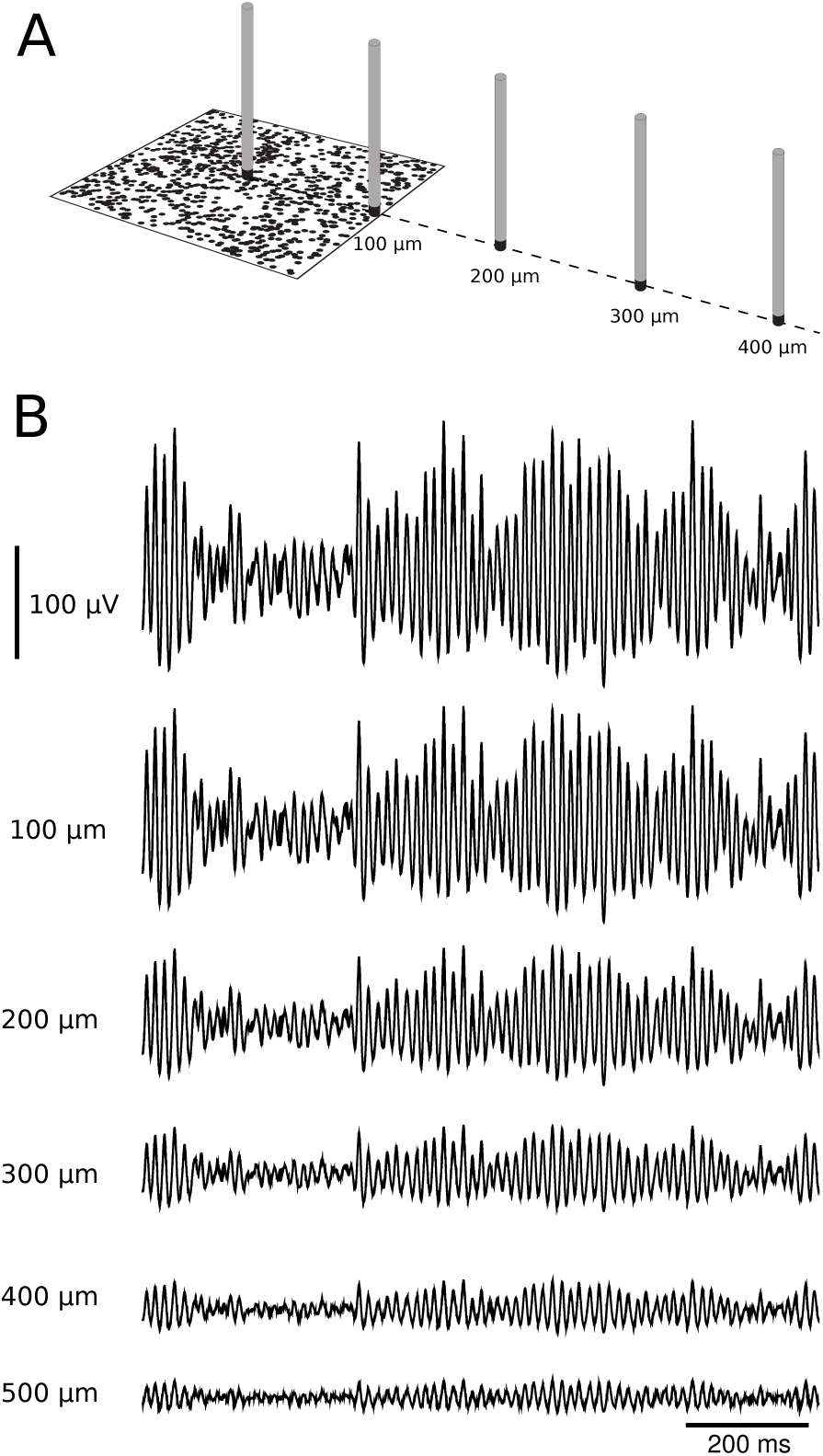
Horizontal distance-dependence of LFP calculated using the kernel method. A. Scheme of the network and the placement of recording sites at different distances from the center of the network. B. LFP calculated (same simulation as in Fig.6) at different distances, as indicated.

In a second example, we used network models capable of generating asynchronous-irregular (AI) or Up/Down state dynamics, which required neurons with spike-frequency adaptation. Fig. 8 illustrates such dynamics as modeled by networks of Adaptive Exponential (AdEx) point neurons (Destexhe, 2009; Zerlaut et al., 2018) (see Methods), as shown in with the associated LFP calculated with the kernel method. A first regime is the asynchronous-irregular state (Fig. 8A), which displays typical LFPs of low-amplitude and noisy aspect, typical of the so-called “desynchronized” dynamics. A second regime is the alternating Up and Down states (Fig. 8B), in which the network produces slow-wave oscillations with higher amplitude LFPs, which was obtained here with strong level of adaptation and additive noise (see Methods). In both cases, the kernel method could simulate the LFP in different cortical depths.

**Figure 8:**
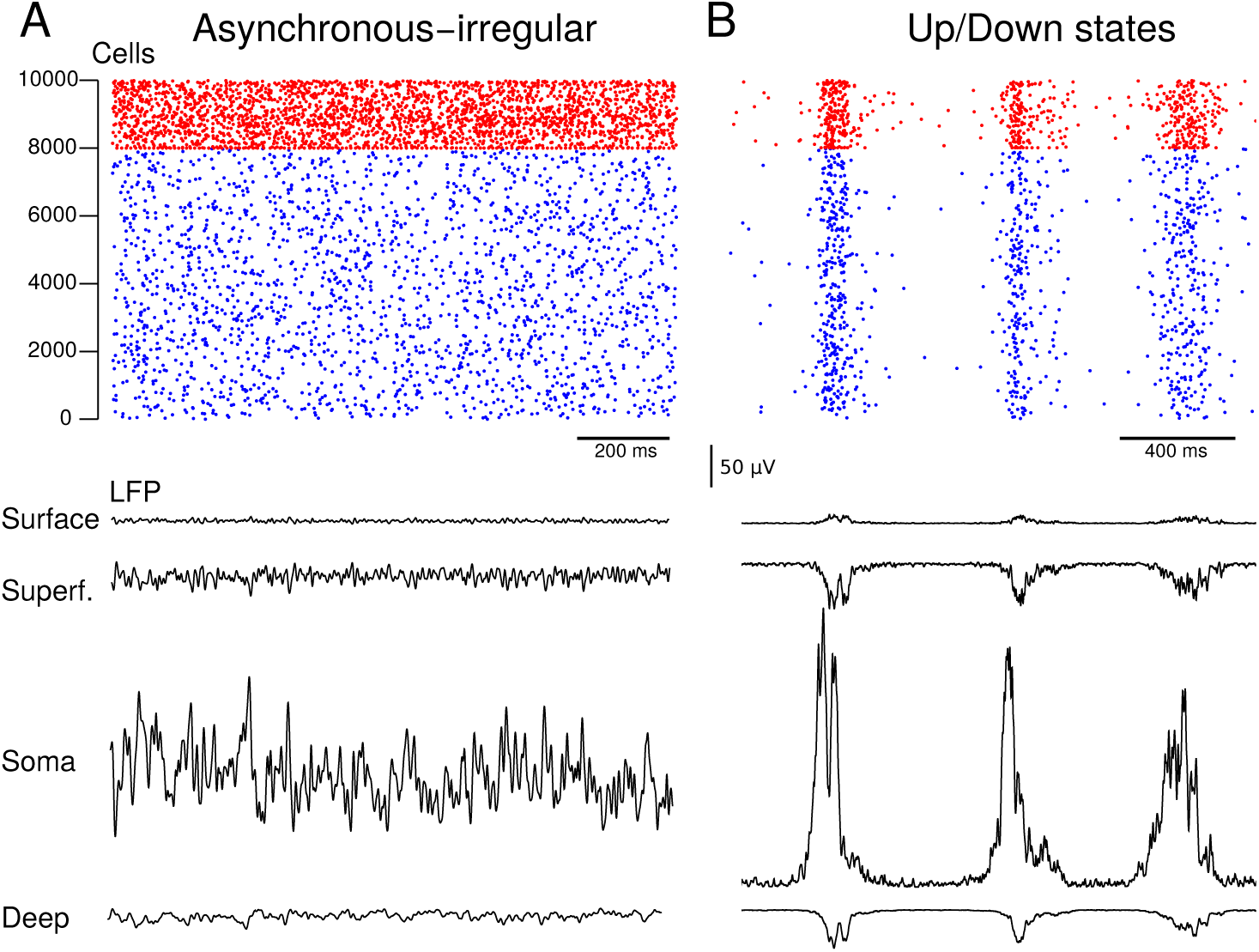
Example of LFP calculated from networks of point neurons in asynchronous or Up/Down states. A. Model of asynchronous-irregular activity in a network of adaptive exponential (AdEx) neurons. The corresponding LFP is calculated and shown identically as in Fig. 6A. The network had 10,000 neurons, 8,000 excitatory (blue) and 2,000 inhibitory (red). B. Same network but for increased adaptation, displaying alternating Up and Down states. The corresponding LFP showed slow wave activity.

To compare the LFP generated by the kernel method to other more classic ways of calculating LFPs, we have considered the method to generate LFPs from synaptic currents (see Methods). As shown in Fig. 9, the LFP was calculated from networks displaying Up and Down states, using the two methods. The LFPs were calculated for surface and depth locations, using the same setting as in Fig. 6. The classic method of LFP generated from synaptic currents (Fig. 9B, gray traces) gave LFPs that were identical from surface and depth, because this method only considers the distance, and the two locations were symmetrical with respect to the network. In contrast, the LFP generated by the kernel method (black traces in Fig. 9B) were different from surface to depth, as described above. Comparing the two methods, the classic method has evidently more high-frequency components, while the kernel method was more smooth. The power spectral density (PSD) calculated from the two models shows this difference explicitly (Fig, 9C). One can also see that the low frequency components were more similar, so the kernel method appears grossly (with some error) as a low-pass filtered version of the classic method based on synaptic currents.

**Figure 9:**
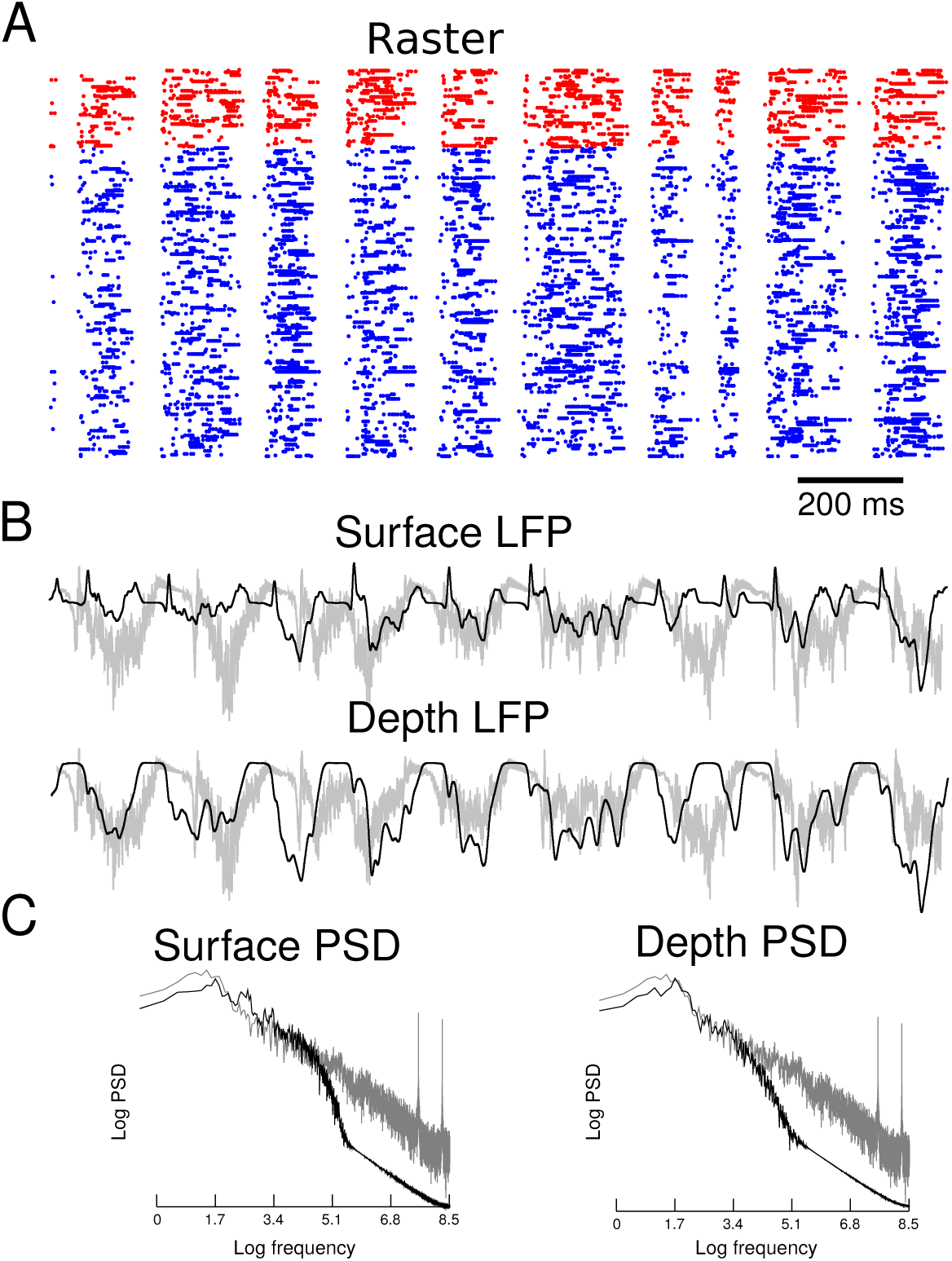
Comparison of the LFP generated by the kernel method to the LFP generated by synaptic currents. A. Raster of spiking activity in a N=5000 network displaying Up and Down states, similar to Fig. 8. B. LFP calculated in surface and depth, using the same scheme as in Fig. 6. The Kernel method (black curves) is compared to the synaptic current method (gray; arbitrary units for amplitude). Note that the gray traces are identical because the LFP generated by the synaptic current method only depends on distance and does not distinguish surface from depth. C. Power spectral density (PSD) calculated from the two models (arbitray units for amplitude).

Finally, we illustrate that the kernel method can be used to calculate the LFPs from multi-layer networks (Fig. 10). As illustrated by the scheme of Fig. 10A, the same network as in Fig. 8B was distributed in three different layers, representing the Supragranular, Granular and Infragranular cortical layers. Taking the same four vertical layers as in Fig. 6, combined for the three networks, leads to six different layers (Surface, Superficial, Supragranular, Granular, Infragranular and Depth). The corresponding LFP calculated in each layer is shown in Fig. 10B. The LFP inverted in superficial and deep layers, as typically found for slow waves between superficial and infragranular layers (Fiáth et al., 2016). It is also consistent with the inversion of the slow wave induced by sensory stimulation (Di, Baumgartner, & Barth, 1990), which also shows this superficial inversion, but in addition a further inversion below infragranular layers, which was also present here (Fig. 8B, Deep).

**Figure 10:**
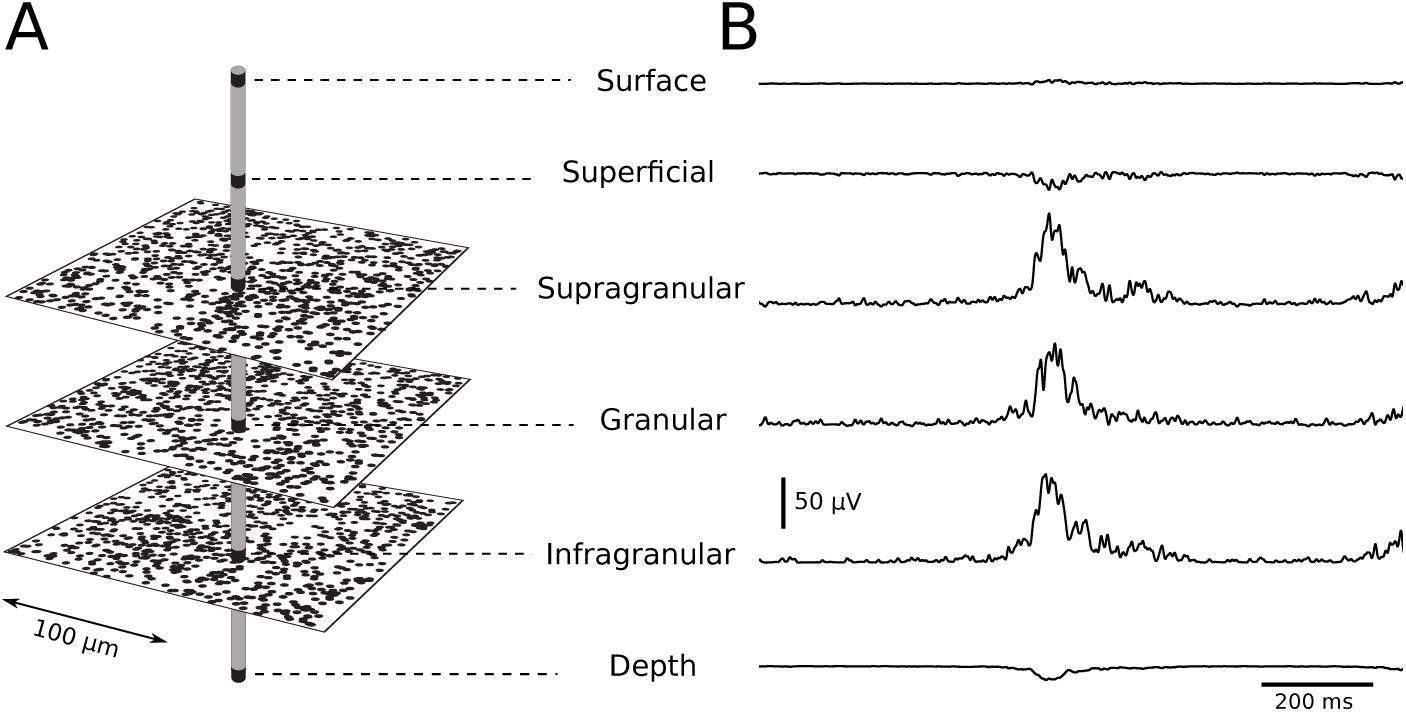
LFP calculated from multilayer networks. A. Three networks similar to Fig. 8B, exhibiting Up/Down state dynamics. The networks are arranged according to three neuronal layers, Supragranular, Granular and Infragranular, as indicated. B. Slow-wave LFPs generated from these networks using the Kernel method. The LFP in 4 layers are generated as in Fig. 8B for each network, and combined, to yield 6 layers, as indicated.

Note that all the examples shown here concerned the cerebral cortex, where the uLFP kernels were measured from experimental data (B. Teleńczuk et al., 2017), but the exact same approach can be followed for any other brain structure.

## 4 Discussion

In this paper, we have proposed a simple method to calculate LFPs from networks of point neurons. We discuss below different aspects of this method, its drawbacks and advantages, and perspectives for future work.

The kernel-based method illustrated here is based on experimentally-measured uLFPs, and is thus dependent on the availability of such measurements. We have used here uLFPs measured in human cerebral cortex, which were obtained in superficial layers (Layer 2-3) (B. Teleńczuk et al., 2017). To complete this dataset, we have used the results from uLFPs calculated from detailed biophysical models (M. Teleńczuk et al., 2020), resulting in the estimated amplitudes displayed in Table 1. As we have illustrated in Fig. 6, this procedure can be used to calculate the LFP in different layers, from network simulations of point neurons.

A first drawback of such a procedure is that we had to use a mix of experimental and computational model data to capture the kernel in different layers. This was done because there is presently no measurement of uLFP in different cortical (or hippocampal) layers, but we anticipate that such data should become available soon given the progress in multielectrode recording techniques, which should release us from using the computational model. When the full data set of uLFPs from all layers, and all cell types will be available, the exact same approach of fitting templates can be followed, and applied to network simulations. Similarly, it may be that the uLFP differs in different cortical regions, due to differences of local connectivity, differences of conductivity, or axonal propagation speed, among other factors. Here again, when the experimental recordings will become available, the method will be refined accordingly.

Another drawback is that the kernel method best applies to (on-going) recurrent activity, because the LFP is exclusively calculated from superthreshold spiking activity. The method does not include contributions such as subthreshold synaptic events (which could be recurrent or evoked), nor the possible contribution of dendritic voltage-dependent ion channels. These different contributions should be estimated by network models of detailed morphologically-reconstructed neurons, where both recurrent and evoked synaptic activity are present.

An important advantage of the present method is the fact that it only relies on the knowledge of cell positions and spike times, which represents a relatively small dataset compared to the knowledge of all membrane currents required by methods to calculate LFPs from biophysical models (Lindén et al., 2014; B. Teleńczuk & Teleńczuk, 2016). As a consequence, the kernel-based method could be applied *a posteriori* to datasets of spike times from network simulations, or even to experimental data, if a sufficiently large number of neurons can be recorded. When the spiking activity will be available from large ensembles of simultaneously-recorded neurons, the kernel-based method could be used to calculate the LFP from spikes, and compare to the recorded LFP, which would constitute a possible test of the consistency of the method.

Another advantage of the kernel-based method is that it does not make any a priori assumption about the conductive or capacitive nature of extracellular media, which is a subject highly discussed in the literature (see (Bedard & Destexhe, 2012; Destexhe & Bedard, 2013) for reviews). Most of today’s procedures to calculate LFPs assume that the extracellular medium is resistive (see for example (Lindén et al., 2014; B. Teleńczuk & Teleńczuk, 2016)), which may result in large errors if it appears that the medium has diffusive or capacitive properties. For example, non-resistive media can exert strong frequency filtering properties which may affect the shape and propagation of LFPs (Bedard & Destexhe, 2012). Another source of frequency filtering is due to the cable properties of the neurons (Pettersen & Einevoll, 2008). In the present method, there is no need to integrate such complex effects, as the method is based on direct recordings of the LFP, so the frequency-filtering, if present, is already taken into account.

The kernel-based method of course does not replace biophysical simulations, which still represent the most accurate way of modeling LFPs. However, such calculations require to have access to the details of the morphology of dendrites, details about the conductivity and other properties of media (as discussed above), and details of all the ionic currents that could influence the LFP. None of such details are needed in the kernel-based method, which calculates LFPs solely from the spiking activity of the neurons. Thus, the kernel method could also be applied to biophysical models, and compared to the LFP generated by standard biophysical methods (Lindén et al., 2014). Such a comparison should be done in future work.

Such as comparison was done previously for the classic method to calculate LFPs from synaptic currents (Mazzoni et al., 2015). It was found that the LFP calculated from a weighted sum of excitatory and inhibitory synaptic currents provides a good approximation of the LFP calculated using morphologically-accurate models. We find here that computing the LFP from synaptic currents of point neurons, which was done in many previous studies (Koch & Segev, 1998; Destexhe, 1998; Mazzoni et al., 2015) is different from the kernel method. This is expected because the kernel method uses a different respective weight of excitation and inhibition, as a function of depth. The method based on synaptic currents only depends on distance, so for instance, it predicts the same LFP for surface and depth if they are at equal distance from the network, as in the example of Fig. 9. The two methods also differ in the high-frequency components of the LFP, but had more similar low-frequency components (Fig. 9C). Thus, the kernel method appears close to a low-pass filtered version of the LFP calculated from synaptic currents. A more in-depth comparison should be done using morphologically-detailed models.

Another method to calculate LFPs from point neurons consists of replaying the membrane currents of the point neurons inside morphologically-accurate models (Hagen et al., 2016). This so-called “hybrid” method can also be used to estimate LFP kernels and use a similar convolution as we used here. However, this approach focuses mostly on the pre-synaptic contributions to the LFP, whereas in the present method, we estimate the LFP from the post-synaptic consequences of axon-propagating action potentials.

Finally, another promising application of the kernel-based method is that it could be applied to population or mean-field models. Since the LFP is obtained by a convolution of the kernel with spiking activity (see Eq. 7), the same approach can be used to convolve the kernel with the density of spiking activity, which is given by mean-field models. This would yield an estimated LFP from mean-field models, which is presently lacking. This also constitutes a very promising direction for future work.

## Acknowledgments

This work was supported by the Centre National de la Recherche Scientifique (CNRS, France), the European Community Future and Emerging Technologies program (Human Brain Project, H2020-720270 and H2020-785907), the ANR PARADOX, and the ICODE excellence network. We thank Eduarda Susin and Matteo di Volo for help with the simulation of the network models.

## Author contributions

BT: Analysis, writing and editing; MT: Analysis, writing and editing; AD: Conceptualization, supervision, analysis, original draft preparation and editing.

